# Should elephants graze or browse? The nutritional and functional consequences of dietary variation in a mixed-feeding megaherbivore

**DOI:** 10.1101/2024.11.13.622959

**Authors:** Hansraj Gautam, Fabio Berzaghi, M Thanikodi, Abhirami Ravichandran, Sheshshayee M. Sreeman, Mahesh Sankaran

## Abstract

Unlike specialist browsers and grazers, the diets of mixed-feeding megaherbivores like elephants are complex and broad, comprising numerous plant species of variable nutritional quality. Here, we revisit an unresolved debate on whether browse is more nutritious than grasses for elephants, as browse is thought to have greater crude protein content. We first analyzed carbon isotopes in 102 fecal samples of Asian elephants to quantify the contribution of browsing and grazing to their diet in Nagarahole National Park, southern India. We show that elephants are predominantly browsers in the grass-scarce Nagarahole forests, in contrast with predominant grazing reported in the nearby grass-abundant savannas of the Nilgiri Biosphere Reserve. We then compared the forage quality of high-browsing and high-grazing diets by analyzing these samples for two proxies of crude protein content (CP): nitrogen content (N%) and Carbon-to-Nitrogen ratio (C:N). Interestingly, high-browsing diets did not have higher N% or lower C:N (proxies for high CP) than low-browsing (high-grazing) diets. To explore the generality of this finding, we analyzed nutritional differences (i.e., CP and fibre content values) between browse and grass across 141 plant species consumed by Asian elephants. We show that woody browse and non-legume browse, which are major components of elephant browse, do not have appreciably higher CP than grasses. Our findings suggest that browsing and grazing broadly have similar nutritional value for such bulk-feeding mixed feeders. Finally, based on the observed habitat-wide variation in browsing, we provide a new framework for assessing how elephants shape woody vegetation across the forests and savannas of Asia, with important implications for conservation and carbon cycling.

## INTRODUCTION

The grazing-browsing spectrum, i.e. variation in the consumption of grass or browse (non-grass) plants, is a key axis along which the foraging ecology of large mammalian herbivores can be differentiated (Jarman, 1974, Codron et al., 2007a, Hempson et al., 2015, Potter et al., 2022). As feeding on grass and browse involves different foraging and digestive mechanisms due to differences in their nutrition, palatability and spatial distribution (Jarman, 1974, Owen-Smith, 1988, Codron et al., 2007b), large herbivores are often classified into grazer, browser, or mixed-feeder (grass-browse mixed consumers) guilds which also map on to their potential impacts on vegetation structure and composition (Owen-Smith, 1988, Hempson et al., 2015, Sabo, 2019, Sitters & Olde Venterink, 2021). The diets of mixed-feeders are particularly interesting and challenging to study since they are complex and broad, comprising hundreds of plant species of varying nutritional quality (Blake, 2002, Sukumar, 2003). This paper examines the diet of Asian elephants (*Elephas maximus*), a mixed-feeding megaherbivore, and revisits an unresolved debate on whether browse is a nutritionally better food type than grasses (Sukumar & Ramesh, 1995, Cerling et al., 1999, Sukumar, 2003 pp. 191-239, Baskaran et al., 2010).

As elephants are hind-gut fermenters, their efficiency of digesting and absorbing nutrients from plant fibers is limited (unlike the ruminants that efficiently absorb nutrients in small intestines after fermentation in the foregut, Owen-Smith, 1988, pp. 78). However, their fast ingesta-passage allows elephants to bulk-feed on a variety of plants for prolonged hours (Blake, 2002, Baskaran et al., 2010); this translates into a generalist diet with massive daily food intake (∼140-200 kg biomass) which compensates for less efficient digestion/absorption (Owen-Smith, 1988). However, all plants are not equal in nutrition (Sukumar, 1990, Codron et al., 2007b, Ganqa et al., 2005, Sitters & Olde Venterink, 2021), and although indiscriminate foraging maximizes biomass consumption, allocating some effort to obtaining high-quality foods helps achieve a more favorable nutrient budget and improved body condition (Owen-Smith & Chafota, 2012, Sukumar, 2003 pp. 191-239).

While the availability of grasses and browse influences what elephants eat (Sukumar, 2003), two compilations of the composition of Asian and African savannah elephant diets indicate that modern elephants predominantly browse (based on carbon isotope signatures in teeth and bone collagen samples, Sukumar & Ramesh, 1995, Cerling et al., 1999 pp. 366), sometimes even in grassy ecosystems (Sukumar, 2003 pp. 191-239). Considering this, Sukumar et al. (Sukumar, 2003 pp. 191-239, Sukumar & Ramesh, 1995) have asserted that browse is nutritionally more important than grasses for elephant growth as 1) it contains higher crude protein content (CP) than grasses (Sukumar, 1990), and 2) proteins containing browse-associated carbon isotopes dominantly contribute to bone collagen in both Asian and African elephants (≥70% proteins came from C3 plants which are mostly browse) (Sukumar & Ramesh, 1992, 1995). Studies on other ungulates also suggest that high-browsing diets have higher CP than grazing-dominated diets (Codron et al., 2007a, Bowman et al., 2010). Sukumar et al. inferred this nutritional importance of browse after finding that browse contributed dominantly to the bone collagen despite contributing equally (as grass) to the food intake by Asian elephants in southern India (Sukumar & Ramesh, 1992, 1995, Sukumar 2003 pp. 203-204; this inference accounted for the enrichment of the heavier isotope due to fractionation during metabolism, Sukumar et al., 1987). Others have cautioned against this inference since elephants were grazers for majority of their recent evolutionary history (Cerling et al., 1999, pp. 369) – with the expansion of C4 grasslands in the Late Miocene, elephants increased C4 grass consumption till Late Pleistocene and still retain the dental features adapted to the abrasive nature of grasses and dust (Saarinen & Lister, 2023). Notably, Baskaran et al. (2010) contested both the importance and predominance of browse vis-à-vis grass in the diet of Asian elephants, and suggest that grass is their principal food while browsing predominates when grass is scarce (see also Baskaran et al. 2024). We attempt to resolve this “browse vs. grass debate” by examining whether browsing is nutritionally better than grazing. Answering this question informs the fundamental ecology of such mixed-feeding megaherbivores (Sukumar, 2003 pp. 191-239, Baskaran et al. 2010, Gautam et al. 2019) and has implications for the management of their habitats (Sukumar & Ramesh, 1995 pp.372, Baskaran et al. 2024 pp. 13-14).

To address this debate, we studied the diet of wild Asian elephants in the heterogeneous Nilgiri Biosphere Reserve (NBR) landscape in southern India. We characterized the grass-browse composition of elephant diets by analyzing carbon isotopes in fecal samples which give a more direct estimate than analyses of bone tissues (for which large adjustments are required, eg. Sukumar & Ramesh, 1992 pp. 538). We collected fecal samples from the tropical deciduous forests of Nagarahole National Park in the NBR. As Nagarahole forests have lower grass abundance than the previously studied grass-abundant mesic savanna habitats in this landscape (Sukumar & Ramesh, 1992, Baskaran et al., 2010, Ahrestani et al., 2012), a study in these forests broadens our understanding of dietary variation in elephant populations. We combined two approaches to address this browse vs. grass debate. First, we analyzed nutrients in the fecal samples from Nagarahole to test the claim that browsing is nutritionally better for elephants than grazing, by quantifying two measures of CP as well as forage quality: nitrogen content (N%) and carbon-to-nitrogen ratio (C:N) (fecal samples are informative of the diets of large herbivores as most of the ingesta comes out undigested which approximates consumed forage, Codron et al., 2005, Leslie et al., 2008). Second, we used two global datasets on plant nutritional quality and plant species consumed by Asian elephants to further test the generality of the claim that browse has better forage quality than grasses. Finally, since dietary variation can shed light on the top-down effects on vegetation (Sabo, 2019), we discuss how the habitat-wide variation in grazing and browsing by elephants can shape their impacts on vegetation and carbon cycling.

## RESULTS

We characterized the grass-browse composition of elephant diet by quantifying carbon-isotope ratios (δ^13^C) in 102 fecal samples collected from three distinct seasons from Nagarahole forests (Methods). The overall mean δ^13^C of dung was -26.05 ‰, indicating that elephants in Nagarahole forests had a heavily browse-dominated diet (≥75% browse), in contrast with grazing-dominated diet observed in the nearby savannas of the same NBR landscape (δ^13^C ∼ -21 to -14 i.e., around ≥50% to ≥90% grass), Ahrestani et al., 2012, Supplement text a). Both the browsing levels and the forage quality of elephant diet declined from wet to dry season (Figure S1, Supplement text b).

### Browse-heavy diets do not have higher crude protein content (CP)

We quantified two proxies of CP in these fecal samples, N% and C:N ratio (Methods) and tested for the effect of diet composition (δ^13^C) on these proxies of CP using linear mixed effects models, while including the effect of season and its interaction with δ^13^C. N% (a positive correlate of CP) was not higher in samples showing browse-heavy diets (i.e. more negative δ^13^C), as evident from the absence of a negative relation between δ^13^C and N% (Table S1, Figure 1a). Instead, a weak positive relation suggests that high-browsing diets were of rather lower forage quality (estimates for δ^13^C’s effect on N%=0.03, Supplement Table S2). Declining browsing levels (i.e., increasing δ^13^C) appeared to lower the N% in wet season as compared to increasing the N% in dry seasons, although this interaction effect was not significant (Table 1a, Table S2). C:N ratio, a negative correlate of forage quality, declined weakly when browsing decreased, indicating that lower-browsing diets were of higher quality (estimate for δ^13^C effect on C:N=-0.04). This relationship varied seasonally (Table S1), with a decline in browsing levels leading to a weak increase in C:N during the wet season but lowering the C:N (i.e. increased forage quality) in the dry season(Figure 2a, Table S2). Overall, the small effect sizes for the main effect of δ^13^C on these proxies of CP imply that the variation in CP was poorly explained by grazing/browsing levels (Table S2). Put simply, higher browsing did not provide higher CP.

**Figure 1.**
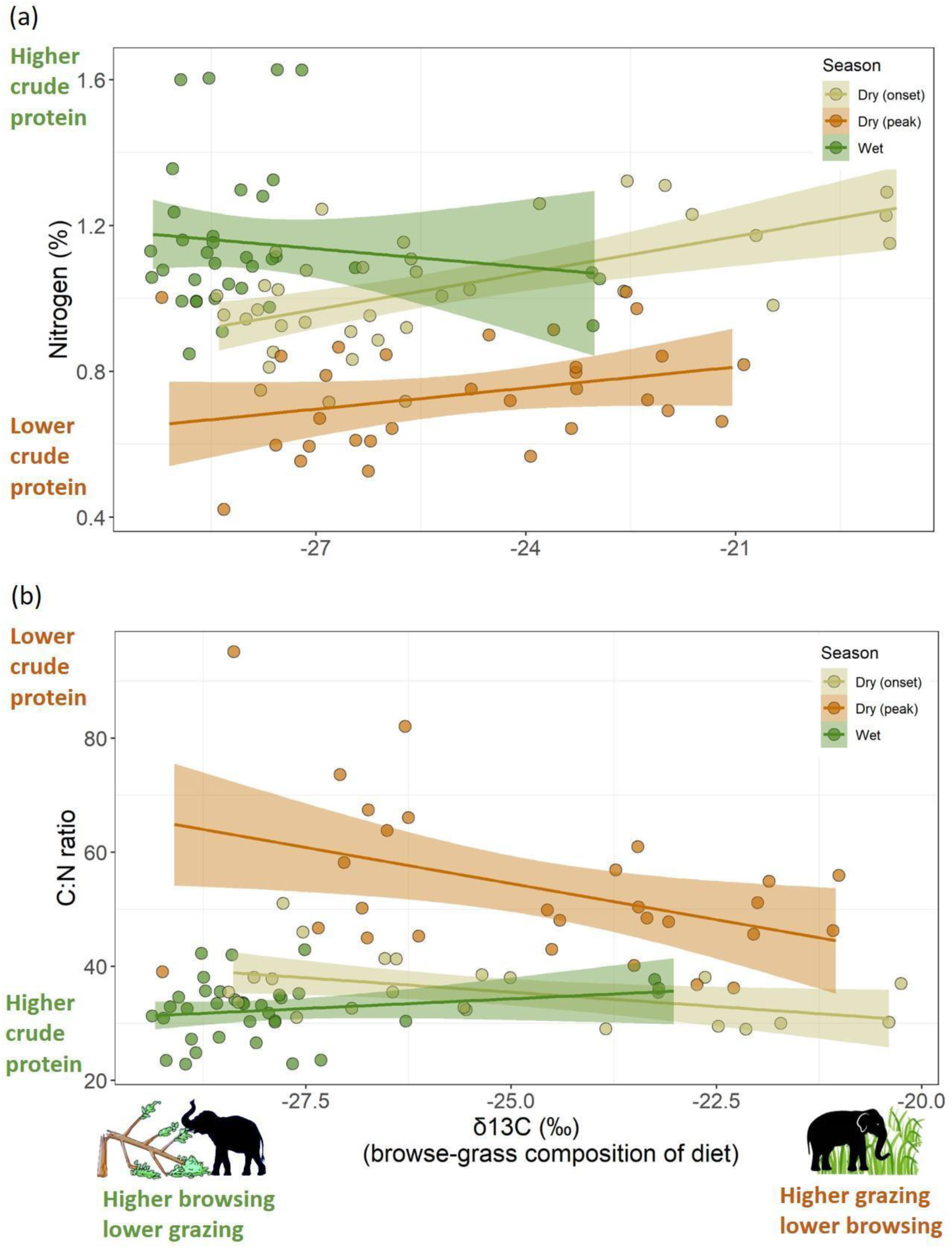
The relationship between grass-browse composition of elephant diet and its nutritional indicators, a) N% and b) C:N ratio. Each circle represents a fecal sample.

**Figure 2.**
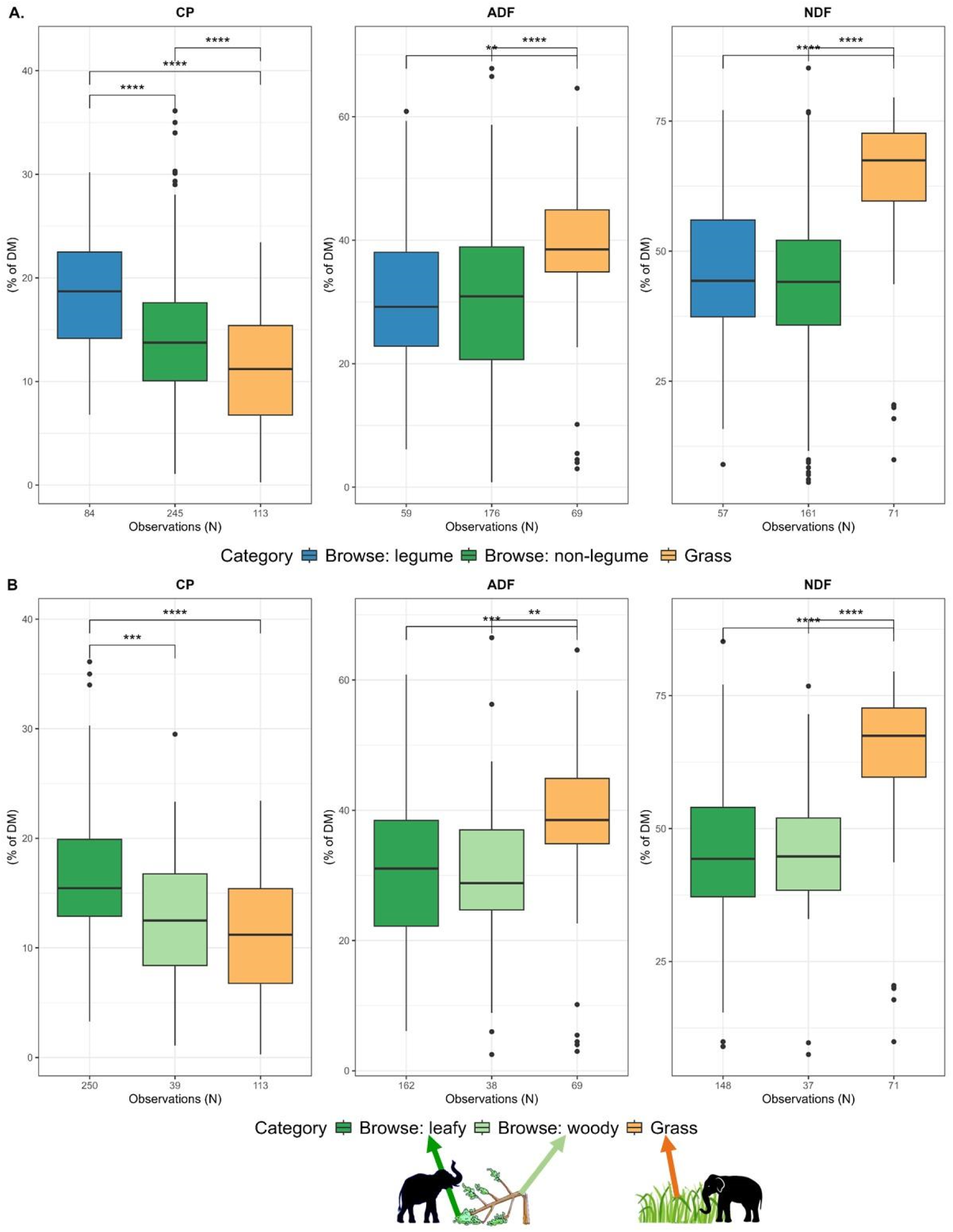
Nutritional values of elephant food species from samples at global extent. **A**. Legume and non-legume browse have higher CP and lower fibre content than grass; **B**. Leafy browse has greater CP and lower fibre content than grass while woody browse has similar CP as grasses. P-values calculated using the t test indicate statistical significance between the mean of the two groups. Significance levels: .P < 0.10; *P <0.05; **P < 0.01; ***P < 0.001; ****P < 0.0001.

### Forage quality of grass and browse food plants

To examine any generalizable differences in the forage quality of browse and grass foods, we analyzed nutritional values (CP, acid detergent fibre (ADF) and neutral detergent fibre NDF) of plants recorded to be consumed by Asian elephants. The list of plant species in Asian elephant’s diet and their nutrient values were extracted from two global databases (Berzaghi & Awasthi, 2022, Berzaghi et al., 2022, Methods). Nutritional data showed that plant species consumed by Asian elephants can also be found outside their current range; to avoid potential geographic bias, we analyzed nutritional data at two spatial extents: one limited to Asia and the other with global extent yielding a larger sample size. At a broader-level categorization of plants (i.e., browse and grass), we found that browse had higher CP and lower fibre content in both Asia-limited and global samples (Figure S2, mean, s.d. and N presented in Table S3-4). However, as we expected browse to be a nutritionally heterogeneous category, we further explored nutritional differences along two sub-categories of browse (Methods). First, we compared grass with legume browse and non-legume browse, as the protein-rich legumes tend to be concentrated in arid habitats (Pellegrini et al., 2016), while Asian elephants live in largely mesic habitats. Legumes clearly had higher CP (18.5% of dry mass) and lower fibre than grass as well as non-legume browse, whereas the CP in non-legume browse (14.3% of dry mass) was only marginally higher than grass (11.6% of dry mass) in the global dataset (Figure 2A, Table S4, Asia-limited results in Figure S3A, Table S3). Second, we compared grass with leafy and woody browse (Methods), as woody browse often dominates elephant browse but may be low in protein. Indeed, only leafy browse had higher CP than grass whereas woody browse did not have significantly higher CP, although it had lower fibre content than grasses (global data: Figure 2B, Table S4; Asia-limited data: Figure S3A, Table S3). Thus, woody browse and non-legume browse, which often dominate the browse component of Asian elephant diet, do not provide appreciably higher CP than grasses.

## DISCUSSION AND CONCLUSION

Here, we revisited the debate on whether browse food plants provide greater protein content and are thus nutritionally more important than grasses for elephants (Sukumar, 2003, Sukumar & Ramesh, 1995, Baskaran et al., 2010). We examined two proximate lines of evidence – (1) the association between carbon isotope ratios (proxies of grazing/browsing levels) and the nutrients in fecal samples indicative of elephant diet, and (2) comparison of the forage quality of browse and grass foods. We found that higher browsing diets (inferred from lower δ13C) did not have appreciably higher CP (inferred from N% and low C:N), which contradicts the presumed nutritional advantage of browse. This trend appears to be generalizable since our analysis of nutritional values of elephant food plants showed that crude protein content in browse, especially woody browse, was not necessarily better than grasses. We also found that browsing levels in elephant diets in Nagarahole forests were much higher than in the savannas of the NBR landscape. Such dietary variation has implications for the impact of these megaherbivores on vegetation.

### Browsing is not nutritionally better than grazing

The similarity in the crude protein content of high-browsing and low-browsing diets (Figure 1) contradicts the presumption that browsing has higher crude-protein value than grazing (Sukumar & Ramesh, 1995, Sukumar 2003). This contradiction is not surprising since elephant browse is a nutritionally heterogeneous category that also contains many low-quality food items. It is worth noting that Sukumar’s inference (2003) of higher protein in browse was based only on leaves (Appendix 1 in Sukumar, 1990) which often represents only a small subset of the elephant diet, especially in the dry season. Woody food items like stems, twigs and bark can constitute as much as leaves, or even dominate the browsing component of elephant diets (eg. 65% of all browse in moist and 37% in dry forests was woody browse in Baskaran, 1998 pp. 96; 66% in Easa, 1999 pp.54, Owen-Smith & Chafota, 2012), but have lower protein than browse leaves (Figure 2). Indeed, in our results on nutrient value of elephant food plants, browse was a nutritionally heterogeneous category, with woody browse having similar CP as grasses and only leafy browse having higher CP (Figure 2), although grasses had the highest fiber content that leads to lower digestibility. Similarly lower CP value of woody browse vis-à-vis leafy browse is also seen in the diet of Africa’s black rhinoceros (Ganqa et al., 2005). Interestingly, CP in leafy browse seems to be higher than in ‘fruit & seed’ although ‘fruit&seed’ were more digestible in our data (Table S3-4). These findings suggest a large nutritional asymmetry along the plant-part axis vis-à-vis the grass-browse axis. Further, while elephant browse as a category did have higher CP (Figure S2, possibly because the data availability was higher for leafy browse Figure 2B), this advantage declined sharply when we excluded legumes (Figure 2) which are usually rare in mesic habitats of Asian elephants. Thus, most of the browse eaten by these bulk-feeding mixed feeders does not provide greater protein than grasses (but see Sitters & Olde Venterink, 2021 for browser vs. grazer guilds), although lower fibre content implies that browse is more digestible than grass. We think that such similarity in the nutrient value of grazing and browsing may be common in other large mammalian herbivores, many of which are mixed feeders (Ahrestani et al. 2016).

Thus, we are faced with two contradictory patterns in the browse vs. grass debate: 1) higher browsing does not imply higher crude protein content(Figure 1 and 2), but 2) browse-associated proteins contribute dominantly to elephant bone collagen (Sukumar & Ramesh, 1995, 1992). Contesting Sukumar’s inference based on δ^13^C analyses of bone collagen, Baskaran et al. (2010) invoked the lack of knowledge of the ranging history of the sampled dead elephants as browse-dominated signatures could arise if elephants used browse-rich habitats. While we concur that ranging history is informative, we note that this browse-dominated signature is very common in collagen of African savannah and Asian elephants (Cerling et al., 1999, Sukumar, 2003). Future studies can reconcile these two contradictory patterns by exploring: a) if carbon from grass and browse gets differentially allocated in body tissues other than collagen, and b) if grasses provide higher instant energy/resources which are utilized immediately and not allocated to body tissues. Future studies can also add insights by replicating our study in other mixed feeders. Simultaneously collecting fecal and plant samples from multiple sites and seasons is desirable, as our global dataset does not control for spatio-temporal variation in CP.

### Functional implications of the dietary variation across forests and savannas

The higher levels of browsing observed in the grass-scarce Nagarahole forests than in the grass-abundant savannas of the NBR landscape suggests that browsing declines with grass abundance (findings from NBR reviewed in Supplement text a). Another contrast between these savannas and forests is the seasonality in grazing/browsing levels: grazing in savannas dominates elephant diet in wet season but declines in dry season (Baskaran et al., 2010, Sukumar & Ramesh, 1992) whereas elephants in Nagarahole forests have a persistently browse-dominated diet throughout the year (Figure S1a, Supplement text b). Such habitat-dependent dietary variation can shape the impact of such mixed-feeding megaherbivores on woody vegetation (Figure 3).

**Figure 3.**
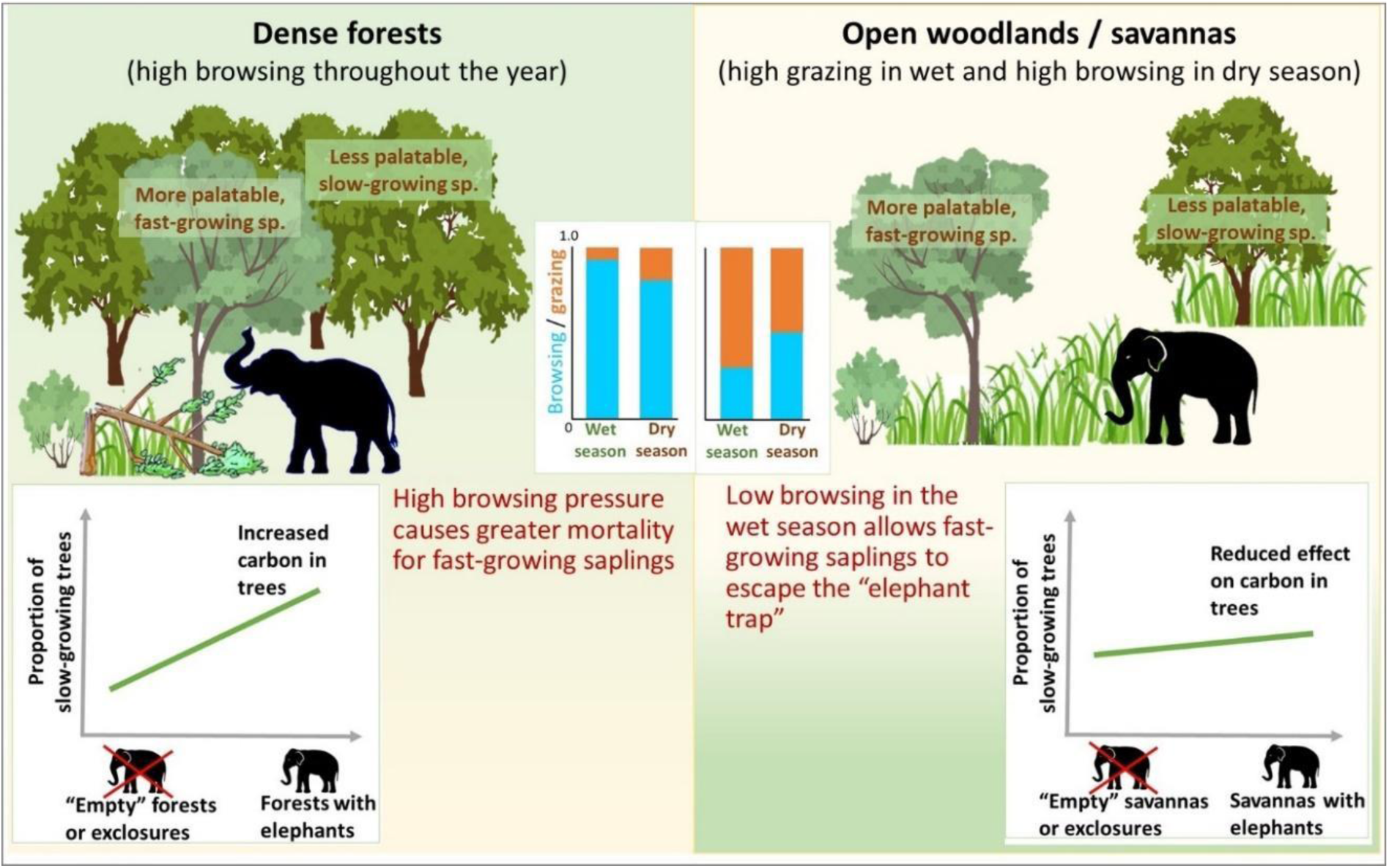
Expected impact of mixed-feeding megaherbivores on woody species composition and aboveground carbon in grass-scarce forests vs. grass-abundant savannas. **Left**: the preferential browsing on fast-growing species with low wood density would increase carbon-dense tree species as compared to the elephant “empty forests” or exclosures, **Right**: high grazing would weaken the browsing effect of elephants on tree carbon as fast-growing, low-carbon trees escape the browse trap. The grazing/browsing levels shown in the inset are qualitative representations based on findings from NBR.

In Figure 3, we present a new framework to assess the impacts of such mixed feeders on woody vegetation and carbon in the savannas and forests of Asia, based on the dietary variation across habitats. In dense forests like Nagarahole, the persistently high browsing levels (due to grass scarcity) create a mortality filter or an “elephant trap” for saplings/recruits (Malhi et al., 2016, Ong et al., 2023). Recently, Berzaghi et al. (2023) showed that African forest elephants seeking high-quality browse cause greater damage to trees of fast-growing species as they are more palatable (due to lower levels of fibres and chemical defenses). Such preferential browsing enhances aboveground carbon storage by shifting tree communities towards slow-growing trees (with higher wood density and carbon storage than faster-growing species) which are less preferred due to higher levels of fibre and defenses (Berzaghi et al., 2023). Given that Asian elephants are browsers throughout the year in forests, they also may potentially promote slow-growing species through the year-round operation of this browsing trap (Figure 3, left). Some empirical support for such impact comes from the preferential browsing causing more damage to tree saplings in early successional forests (dominated by fast-growing species) than in mature forests in Malaysia (eg. Ong et al., 2023). In contrast, in the savannas, some fast-growing tree recruits may escape this mortality filter in the productive season when elephants primarily graze. Such seasonal relaxation of tree mortality may limit the ability of elephants to shift woody community composition towards slow-growing species in savannas (vis-a-viz forests). In other words, their role in such aboveground carbon sequestration is expected to be lower in mesic savannas than forests (flatter slope in Figure 3, right). Secondly, such tree mortality may be critical for the emergence of canopy gaps in forests inhabited by elephants, thus enhancing plant diversity by promoting favorable sunlight niches for shade-intolerant species (Terborgh et al., 2016). Thirdly, these destructive foragers leave behind trails of high disturbance (Blake, 2002), thus promoting heterogeneous microclimates for a diverse understory, through woody mortality in forests and grass removal in savannas. Such diet-mediated impacts of mixed-feeders on vegetation can be tested through observations of foraging (eg. Ong et al., 2023), exclosure experiments or dynamic vegetation models (Berzaghi et al., 2019), and could refine the recent emphasis on the global role of such megaherbivores in shaping vegetation and carbon dynamics.

Our results clarifying the fundamental ecology of this mixed feeder have implications for habitat management. In light of the similar nutritional value of elephant browse and grass, we suggest a moderation in the emphasis on promoting either browse (Sukumar & Ramesh, 1995 pp.372) or grass (Baskaran et al. 2024 pp. 13-14) in Asian elephant habitats. The question of which habitats or mosaics provide nutritionally better forage and thus can better support elephants needs to be revisited, especially because diet varies widely with vegetation composition. Replicating our study in multiple sites (and extending it to other nutrition variables) can be a useful approach to assess forage quality, including in other mixed-feeding large herbivores.

## MATERIALS AND METHODS

### Study area

The study area, Nagarahole National Park (approx. 11.85°–12.26° N, 76.00°– 76.28° E), is part of the Nilgiri Biosphere Reserve landscape (NBR), which (along with Eastern Ghats) supports the world’s largest contiguous population of Asian elephants. Dry deciduous, moist deciduous and teak plantations are the major forest types in Nagarahole and elephants have access to both grasses and browse plants. Grass abundance in these two forest types in Nagarahole (avg. 202 g/m2, Gautam et al., 2017) is less than 1/4th of that from the dry deciduous (921 g/m2) and less than half of that in the dry thorn savanna (524 g/m2) habitats in NBR; predominant grazing was observed by Baskaran et al. (2010) in the latter two habitats. In this paper, we refer to the latter two habitats as savannas due to their open canopy and understory having abundant grasses, following Sankaran & Ratnam (2013). Due to grass scarcity in Nagarahole forests, we expected higher browsing and lower grazing in these forests compared to savannas. To study elephant diets, we adopted the widely used method of studying fecal samples which are reliably informative of diets at short-term scales (1-5 days for elephants), as majority of ingesta passes as undigested material in the faeces of large herbivores and thus reflects the consumed forage (Codron et al., 2005, Leslie et al., 2008, Fernando et al., 2016).

### Sample collection and processing

We (HG and TM) collected elephant dung samples around three different time points spaced at an interval of three months, i.e. 1 October 2022, 1st January and 1st April in 2023. These correspond to the distinct seasons i.e., namely wet season, onset of the dry season, and peak dry season, respectively. We temporally concentrated our sampling such that ∼90% samples (91/102) were collected during a ± 2- week window around these dates. To minimize the confounding effect of age class on diet (Sukumar & Ramesh, 1992), we only collected dung samples with large bolus size matching that of adults. Upon finding fresh dung samples (a few hours old or from overnight), we broke open one dung bolus to collect 5-6 chunks of dung from different parts (around 200- 300 grams of fresh weight), while excluding the outer slimy layer. We assigned an ID to each sample and noted its GPS location. Further, in case we collected multiple dung samples which were within 50m of each other, and thus could belong to the same foraging group, we assigned a common cluster ID to these potentially non-independent samples. Samples were air-dried in the sun to the extent possible, and then transported to the lab in thick-paper bags. We dried them further in a hot-air oven at ∼60° C for ∼60 hours to remove any moisture. Next, we ground these samples to obtain a fine powder for laboratory analyses. We thoroughly mixed the contents of each sample before grinding, so that the powder obtained was representative of the collected dung.

### Quantifying grazing/browsing levels in elephant diet

To quantify the grass-browse composition of diet, we performed mass spectrometry analyses of fecal samples to obtain δ^13^C i.e., the ratio of the heavy ^13^C and light ^12^C stable carbon isotopes (analyses carried out by MS at the National Stable Isotopes Facility at the University of Agricultural Sciences, Bangalore). We quantified the δ^13^C for fecal samples with respect to the reference standard (Vienna Pee Dee Belemite) as:

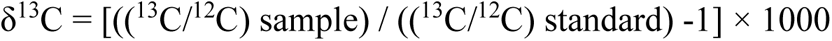

Here, ^13^C and ^12^C represent masses of the two carbon isotopes measured in the sample and the standard.

C3 plants are more depleted in the heavier isotope and thus yield lower (more negative) δ^13^C values. For instance, C3 browse plants in this region have average δ^13^C value of -27.2 ‰ while C4 grasses have average δ^13^C = -12.8 ‰ (Sukumar et al., 1987). When a mixture of C3 and C4 plants (eg. mixed diet) is analyzed, a range of intermediate values of δ^13^C values can be obtained, depending on which type of plants dominate the plant mixture. Controlled feeding experiments on ungulates show very minor differences in δ^13^C between actual forage consumed and faeces, regardless of whether the given forage was dominated by C3 or C4 (consistent difference of around -0.9 ‰, Codron et al., 2005). Thus, the δ^13^C analysis of faeces reliably quantifies diet composition. Here, we qualitatively inferred ∼ 50% contribution of C4 grasses and browse to elephant diet when δ^13^C was around -20, following Codron et al. (2006), although exact composition should ideally be modelled from mixtures of local vegetation.

### Quantifying nutrients in diet

To study nutritional quality of diet, we (HG and AR) examined two metrics in fecal samples: Nitrogen content (N%) and Carbon-to-Nitrogen ratio (C:N), which are positive and negative correlates of diet quality, respectively (Codron et al., 2007, Leslie et al., 2008). N% approximates CP by a factor of ∼ 1/6.25 while C:N is inversely related to CP. We quantified the fecal N% and C% (to obtain C:N ratio) using the Leco TrueSpec CN analyzer at NCBS, Bangalore, using EDTA LCRM® as the reference standard.

### Processing data on the nutritional quality of grass and browse plants eaten by elephants

To test the generality of the claim that browse is more nutritious than grasses, we collated nutritional values of elephant food plants using two global databases: 1) MegaFeed, a database of food plants of extant Proboscidea species, from which we extracted the list of recorded food plants of Asian elephants (Berzaghi & Awasthi, 2022), and 2) PNuts, a global database of plant nutritional values for different type of plants and plant parts or organs (Berzaghi et al., 2022). PNuts has been used in a previous study on nutritional values of forest elephant foods (Berzaghi et al. 2023). PNuts provided values of CP, acid detergent fiber (ADF) and neutral detergent fibre (NDF) for plant species consumed by Asian elephants as indicated in MegaFeed (complete list of plant species and nutritional values in SI Data S1). Plant species names were updated and homogenized following the taxonomy in World Flora Online (https://wfoplantlist.org/). ‘Liana’,’Tree’,’Shrub’,’Legume’,’Herb’ in the ‘Plant_type’ column in Data S1 were classified as “browse”. Some plants consumed by elephants can also be found outside their range, because they are either plants with rather large distribution, or species that have been introduced. Plant nutritional values are determined by plant type but also by their growing environmental conditions (Berzaghi et al. 2022). To avoid potential bias given by growing conditions outside elephant’s range, we analyzed nutritional data both for Asia only and the global dataset.

Fibers were analyzed in addition to protein because low values of ADF and NDF indicate higher digestibility of forage thus higher assimilation of protein and other nutrients. We first explored nutritional differences between a) grasses and browse. We then explored different subcategories of browse for comparison of b) grass with non-legume browse and legume browse, since legumes are rare in the mesic habitats of Asian elephants, and comparison of c) grass with subcategories of browse plant parts (leafy browse and woody browse that included bark, branch, stem, etc.; fruit&seed was excluded from analyses due to small sample size but its values are reported in Table S2-3) since a lot of browse eaten by elephants includes woody tissues (Baskaran, 1998 pp. 96, Easa, 1999 pp. 54) which may be low-quality foods.

### Statistical analyses

First, to test if grass-browse composition predicts crude protein content, we separately modeled N% and C:N in elephant fecal samples using LMMs (lmer function in lmerTest package Kuznetsova et al. 2017), by including δ^13^C, season and their interaction as fixed-effects predictors and sample cluster as random effect. The δ^13^C × season interaction would allow the relationship (slope) between δ^13^C and diet quality to vary in different seasons. In these two LMMs, we modelled the intercept term by limiting δ^13^C (continuous predictor) to its smallest negative value in the dataset (rather than 0 which is an unrealistic δ^13^C value for these plants). To infer the significance of effects, we obtained ANOVA table using anova function with Type III SS appropriate for unbalanced designs (obtained using Wald distribution). We inferred the effect size or slope from the estimates of fixed-effects in the LMMs. We log-transformed C:N to reduce heteroscedasticity and skew in the distribution of residuals in these LMMs. We limit our inferences regarding grazing to only C4 grass consumption because Nagarahole forests also have C3 grasses which have δ^13^C values similar to browse. This does not alter our expectation of higher N% and lower C:N in dung with higher consumption of C3 plants (mostly browse) compared to C4 grasses, since C3 grasses also have higher CP than C4 grasses (Barbehenn et al., 2004). Second, we analyzed data from the two global databases mentioned above to further examine any generalizable nutritional differences between browse and grass. We analyzed plant nutritional values (CP, ADF and NDF) in Asian elephant food plants, along the three categorizations mentioned above. We used pairwise t-tests in R to test statistical differences between the means of different categories: a) grass and browse, b) grass, legume browse and non-legume browse, and c) grass, leafy browse, woody browse. We did these analyses with plant nutrient data coming from two spatial extents: Asia only and global extent.

## ACKNOWLEDGMENTS

We are grateful to the Principal Chief Conservator of Forests, Karnataka Forest Department, the office of the Conservator of Forests and Director, Rajiv Gandhi (Nagarahole) National Park, Hunsur, and other forest officials for field research permits. We especially thank Prof. T.N.C. Vidya and the Kabini Elephant Project for the generous support with logistics during the entire fieldwork as well as for the collaboration under which the fieldwork was carried out. Anvitha S. and Jabili Chowdary helped in collecting some of the dung samples. We thank Ms. Sushma and the National Facility for Stable Isotope Studies at the University of Agricultural Sciences, GKVK, Bengaluru, for δ^13^C analyses. We are thankful to our field assistants Pramod, Krishna and Shankar for assisting with fieldwork and ensuring our safety. We thank Rohit Subedar, Manaswi Raghurama and other members of BEER lab for feedback. HG analyzed and wrote a part of this manuscript while working at the University of Turku.

## REFERENCES

1. Ahrestani, F. S., Heitkönig, I. M. A., & Prins, H. H. T. (2012). Diet and habitat-niche relationships within an assemblage of large herbivores in a seasonal tropical forest. Journal of Tropical Ecology, 28(4), 385–394. 10.1017/S0266467412000302

2. Ahrestani, F. S., Heitkönig, I. M. A., Matsubayashi, H., & Prins, H. H. T. (2016). Grazing and Browsing by Large Herbivores in South and Southeast Asia. In The Ecology of Large Herbivores in South and Southeast Asia (p. 99). Springer.

3. Barbehenn, R. V., Chen, Z., Karowe, D. N., & Spickard, A. (2004). C 3 grasses have higher nutritional quality than C 4 grasses under ambient and elevated atmospheric CO 2. Global Change Biology, 10(9), 1565–1575. 10.1111/j.1365-2486.2004.00833.x

4. Baskaran, N. (1998). Ranging and resource utilization by asian Elephant (Elephas maximus Linnaeus) in Nilgiri Biosphere Reserve South India. Bharathidasan University.

5. Baskaran, N., Balasubramanian, M., Swaminathan, S., & Desai, A. A. (2010). Feeding ecology of the Asian elephant Elephas maximus Linnaeus in the Nilgiri Biosphere Reserve, southern India. Journal of the Bombay Natural History Society, 107(1), 3–13.

6. Baskaran, N., Sathishkumar, S., Vanitha, V., Arjun, M., Keerthi, P., & Bandhala, N. G. (2024). Unveiling the Hidden Causes: Identifying the Drivers of Human–Elephant Conflict in Nilgiri Biosphere Reserve, Western Ghats, Southern India. Animals, 14(22), 3193.

7. Gautam, H., Arulmalar, E., Kulkarni, M. R., & Vidya, T. N. C. (2019). NDVI is not reliable as a surrogate of forage abundance for a large herbivore in tropical forest habitat. Biotropica, 51(3), 443–456.

8. Berzaghi, F., & Awasthi, B. (2022). MegaFeed: Global database of megaherbivores’ feeding preferences. 10.1101/2022.09.23.509174

9. Berzaghi, F., Bretagnolle, F., Durand-Bessart, C., & Blake, S. (2023). Megaherbivores modify forest structure and increase carbon stocks through multiple pathways. Proceedings of the National Academy of Sciences, 120(5), e2201832120. 10.1073/pnas.2201832120

10. Berzaghi, F., Bretagnolle, F., Ratshikombo, Z., & Abdallah, A. B. (2022). PNuts: Global spatio-temporal database of plant nutritional properties. 10.1101/2022.08.29.505708

11. Berzaghi, F., Longo, M., Ciais, P., Blake, S., Bretagnolle, F., Vieira, S., Scaranello, M., Scarascia-Mugnozza, G., & Doughty, C. E. (2019). Carbon stocks in central African forests enhanced by elephant disturbance. Nature Geoscience, 12(9), 725–729. 10.1038/s41561-019-0395-6

12. Blake, S. (2002). The ecology of forest elephant distribution and its implications for conservation. University of Edinburgh.

13. Cerling, T. E., Harris, J. M., & Leakey, M. G. (1999). Browsing and grazing in elephants: The isotope record of modern and fossil proboscideans. Oecologia, 120(3), 364–374. 10.1007/s004420050869

14. Codron, D., Codron, J., Sponheimer, M., Lee-Thorp, J. A., Robinson, T., Grant, C. C., & De Ruiter, D. (2005). Assessing diet in savanna herbivores using stable carbon isotope ratios of faeces. Koedoe, 48(1), 115–124. 10.4102/koedoe.v48i1.170

15. Codron, D., Lee-Thorp, J. A., Sponheimer, M., & Codron, J. (2007). Nutritional content of savanna plant foods: Implications for browser/grazer models of ungulate diversification. European Journal of Wildlife Research, 53(2), 100–111. 10.1007/s10344-006-0071-1

16. Codron, D., Lee-Thorp, J. A., Sponheimer, M., Codron, J., De Ruiter, D., & Brink, J. S. (2007). Significance of diet type and diet quality for ecological diversity of African ungulates. Journal of Animal Ecology, 76(3), 526–537. 10.1111/j.1365-2656.2007.01222.x

17. Codron, J., Lee-Thorp, J. A., Sponheimer, M., Codron, D., Grant, R. C., & De Ruiter, D. J. (2006). Elephant (Loxodonta Africana) diets in Kruger National Park, South Africa: Spatial and Landscape Differences. Journal of Mammalogy, 87(1), 27–34. 10.1644/05-MAMM-A-017R1.1

18. Easa, P. (1999). Status, habitat utilization and movement pattern of large mammals in Wayanad Wildlife Sanctuary. Kerala Forest Research Institute, Peechi.

19. Fernando, P., Jayasinghe, L. K. A., Rahula Perera, R. A., Weeratunga, V., Kotagama, S. W., & Pastorini J. (2016). Diet component estimation in Asian elephants by microhistological faecal analysis. Gajah, 44, 23–29.

20. Ganqa, N. M., Scogings, P. F., & Raats, J. G. (2005). Diet selection and forage quality factors affecting woody plant selection by black rhinoceros in the Great Fish River Reserve, South Africa. South African Journal of Wildlife Research, 35(1), 77–83.

21. Gautam, H., Potdar, G. G., & Vidya, T. N. C. (2017). Using visual estimation of cover for rapid assessment of graminoid abundance in forest and grassland habitats in studies of animal foraging. Phytocoenologia, 47(4), 315–327. 10.1127/phyto/2017/0146

22. Hempson, G. P., Archibald, S., & Bond, W. J. (2015). A continent-wide assessment of the form and intensity of large mammal herbivory in Africa. Science, 350(6264), 1056–1061. 10.1126/science.aac7978

23. Jarman, P. (1974). The social organisation of antelope in relation to their ecology. Behaviour, 48(1–4), 215–267. Kuznetsova, A., Brockhoff, P.B., & Christensen, R.H.B. (2017). lmerTest Package: Tests in Linear Mixed Effects Models. Journal of Statistical Software, 82(13), 1–26 doi.org/10.18637/jss.v082.i13

24. Leslie, D. M., Bowyer, R. T., & Jenks, J. A. (2008). Facts From Feces: Nitrogen Still Measures Up as a Nutritional Index for Mammalian Herbivores. The Journal of Wildlife Management, 72(6), 1420–1433. 10.2193/2007-404

25. Kuznetsova, A., Brockhoff, P.B., & Christensen, R.H.B. (2017). lmerTest Package: Tests in Linear Mixed Effects Models. Journal of Statistical Software, 82(13), 1–26 doi.org/10.18637/jss.v082.i13

26. Malhi, Y., Doughty, C. E., Galetti, M., Smith, F. A., Svenning, J.-C., & Terborgh, J. W. (2016). Megafauna and ecosystem function from the Pleistocene to the Anthropocene. Proceedings of the National Academy of Sciences, 113(4), 838–846. 10.1073/pnas.1502540113

27. Ong, L., Tan, W. H., Davenport, L. C., McConkey, K. R., Amin, M. K. A. bin M., Campos-Arceiz, A., & Terborgh, J. (2023). Asian elephants as ecological filters in Sundaic forests. Frontiers in Forests and Global Change, 6, 1143633.

28. Owen-Smith, N., & Chafota, J. (2012). Selective feeding by a megaherbivore, the African elephant ( Loxodonta africana). Journal of Mammalogy, 93(3), 698–705. 10.1644/11-MAMM-A-350.1

29. Owen-Smith, R. N. (1988). Megaherbivores: The Influence of Very Large Body Size on Ecology. Cambridge University Press.

30. Pellegrini, A. F. A., Staver, A. C., Hedin, L. O., Charles-Dominique, T., & Tourgee, A. (2016). Aridity, not fire, favors nitrogen-fixing plants across tropical savanna and forest biomes. Ecology, 97(9), 2177–2183.

31. Potter, A. B., Hutchinson, M. C., Pansu, J., Wursten, B., Long, R. A., Levine, J. M., & Pringle, R. M. (2022). Mechanisms of dietary resource partitioning in large-herbivore assemblages: A plant-trait-based approach. Journal of Ecology, 110(4), 817–832. 10.1111/1365-2745.13843

32. Sabo, A. E. (2019). Impacts of Browsing and Grazing Ungulates on Plant Characteristics and Dynamics. In The Ecology of Browsing and Grazing II (pp. 259–276). Springer International Publishing.

33. Sankaran, M., & Ratnam, J. (2013). African and Asian Savannas. In Encyclopedias of Biodiversity (2nd ed., Vol. 1, pp. 58–74). Elsevier Press. 10.1016/B978-0-12-384719-5.00355-5

34. Sitters, J., & Olde Venterink, H. (2021). Herbivore dung stoichiometry drives competition between savanna trees and grasses. Journal of Ecology, 109(5), 2095–2106. 10.1111/1365-2745.13623

35. Sukumar, R. (1990). Ecology of the Asian Elephant in Southern India. II. Feeding Habits and Crop Raiding Patterns. Journal of Tropical Ecology, 6(1), 33–53.

36. Sukumar, R. (2003). The living elephants: Evolutionary ecology, behavior, and conservation. Oxford University Press.

37. Sukumar, R., Bhattacharya, S. K., & Krishnamurthy, R. V. (1987). Carbon isotopic evidence for different feeding patterns in an Asian elephant population. Current Science, 56(1), 11–14.

38. Sukumar, R., & Ramesh, R. (1992). Stable carbon isotope ratios in Asian elephant collagen: Implications for dietary studies. Oecologia, 91(4), 536–539. 10.1007/BF00650328

39. Sukumar, R., & Ramesh, R. (1995). Elephant foraging: Is browse or grass more important? In Daniel, J.C. & H.S. Datye (Eds): A Week with Elephants. (pp. 368–374). Bombay Natural History Society, Oxford University Press.

