## Supplementary Material for "Should elephants graze or browse? The nutritional and functional consequences of dietary variation in a mixed-feeding megaherbivore"

**This PDF file includes:**

Supplement text a and b

Figures S1 to S3

Tables S1 to S4

SI References

### SUPPLEMENT TEXT

#### Supplement text a. Habitat-wide differences in the levels of grazing/browsing by elephants

A qualitative comparison of the elephant diet in the forest and savanna habitats of the NBR landscape suggests that diet composition varies with vegetation, especially grass abundance. Our study shows a browse-dominated diet (≥75% browse, based on δ13C <-26‰) in Nagarahole forests where grass abundance is low. In contrast, in the savannas of Mudumalai and Bandipur where grasses are abundant (Baskaran et al., 2010), Ahrestani et al., (2012) found higher δ13C (monthly avg. δ13C ~ -21 to -14), indicating that grazing either matched or exceeded browsing (≤50% was browse). This trend is consistent with observations of collared elephants by Baskaran et al. (2010) who reported that while grazing was high overall (80%), it declined from 91% in tall-grass deciduous habitats (which are technically mesic savannas due to grass-dominated understory and open canopy) to 74% in dry thorn savannas where grass was relatively less abundant, and declined further to 54% in moist deciduous forests with lower grass availability. Finally, Sukumar's (1990) study around Sathyamangalam reported high browsing in short-grass habitats and high grazing in tall-grass habitats or savannas. As browse is abundant in both forests and savanna habitats of NBR, the variation in grazing/browsing levels appears to be contingent on the availability of grasses rather than browse, although detailed measurements would be needed to say this confidently. This cross-habitat trend appears to validate the preference for grasses as suggested by Baskaran et al. (2010, pp. 10), contrary to Sukumar et al.'s suggestion that browse is the more preferred and nutritionally important food type and thus is the limiting factor for elephants (Sukumar & Ramesh, 1995, Sukumar, 2003). Future studies that quantify diet composition in multiple locations along a gradient of grass/browse abundance are needed to examine whether this is a robust trend. While our analyses of nutrient values in grass and browse suggests a similarity in the crude protein value of grass and browse in elephant diet (Figure 2 in main text), the higher digestibility of browse should also be considered a key factor while differentiating the nutritional value along the grazing-browsing spectrum.

**Supplement text b. On seasonal variation in diet composition and quality**

There were seasonal shifts in both diet composition and quality. Elephant diet was heavily browse-dominated in the wet season (δ13C=-27.89 corresponding to >90% diet being browse, see Figure S1a), while grazing seemed to increase marginally in the dry seasons although the diet was still browse-dominated (peak dry season δ13C= -24.84 i.e., >70% was browse). In contrast with this persistent browse-dominated diet in both wet and dry season, elephants in the savannas of NBR shift their grazing-dominated diet in the other direction: grazing declines from wet to dry seasons in the mesic savannas of NBR referred to as dry deciduous (grazing declined from 95% to 85%) and dry thorn habitats (grazing declined from 88% to 53%) in Baskaran et al. (2010) and this shift may relate to seasonal shifts in protein content in grasses as they mature from wet to dry season.

We found more crude protein (high N% and low C:N ratio) in dung during the wet season when browsing was highest, than in dry season when browsing reduced. The fecal N% declined from 1.14% in wet season to 1.02% at the onset of dry season and to 0.71% in the peak dry season. Similarly, C:N ratio increased from 32.2 in wet season to 36.6 at the onset and increased further to 54.7 in the peak dry season, indicating worsening diet quality from wet to dry seasons. The seasonal changes in diet quality tended to be sharper than the changes in grass-browse composition (N%: 10.5% change from wet to the onset of dry season, 30.4% change between the onset and peak of dry seasons; C:N: 13.7% and 49.5%, respectively; δ13C: 9.3%, 1.8%, respectively) (Figure S1). This seasonal decline in CP in diet is consistent with the decline in browsing and increase in C4 grass consumption in the dry seasons.

This trend of diet quality declining with seasonal decline in browsing (Figure S1) may appear to be consistent with Sukumar's suggestion (2003) that browsing is more important than grazing.

However, we caution against inferring that higher browsing provided more protein as season is a confounding variable in our dataset. For example, this pattern may arise from seasonal variation in nutrients due to phenology, like in Mudumalai where foliar nitrogen in herbaceous vegetation declined from wet to dry season (Ahrestani et al., 2011). Further, diet quality could also vary if elephants were eating different browse items in wet and dry seasons based on availability; for example, elephants bulk-feed on leafy herbs like *Globba maratina* and Zingiber sp. that are abundant in wet season but unavailable in dry season (HG personal observations 2011-2016). Moreover, a seasonal switch to low-quality woody items under scarcity of protein-rich foliage could also reduce nitrogen intake. Future studies that simultaneously measure nutrients in the standing vegetation and in elephant diet can provide insights about the sources of such seasonal variation in nutrient intake. DNA-metabarcoding may provide a more detailed taxonomic characterization of elephant diet, in contrast with the broad grazing-browsing spectrum inferred from carbon isotope analyses conducted by us.

**SUPPLEMENT FIGURES**

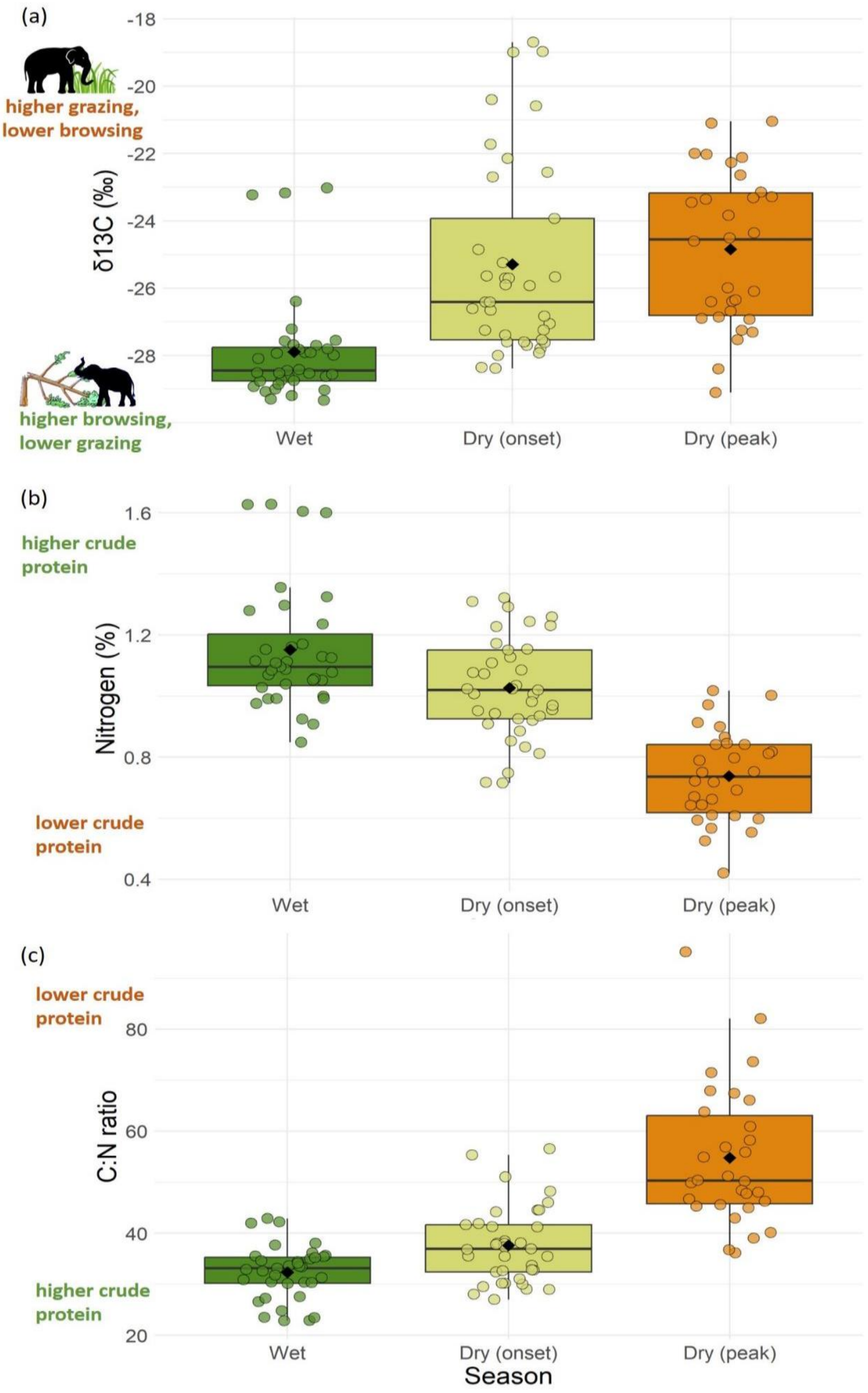


**Figure S1:** Seasonal variation in grass-browse composition and nutritional quality of elephant diet inferred from fecal samples from wet, early dry and peak dry season. As the dry season progresses **a)** consumption of C4 grasses increases, **b)** nitrogen content declines, **b)** C:N ratio increases. Each circle represents a fecal sample, while black diamonds represent seasonal means and the horizontal line represents the median.

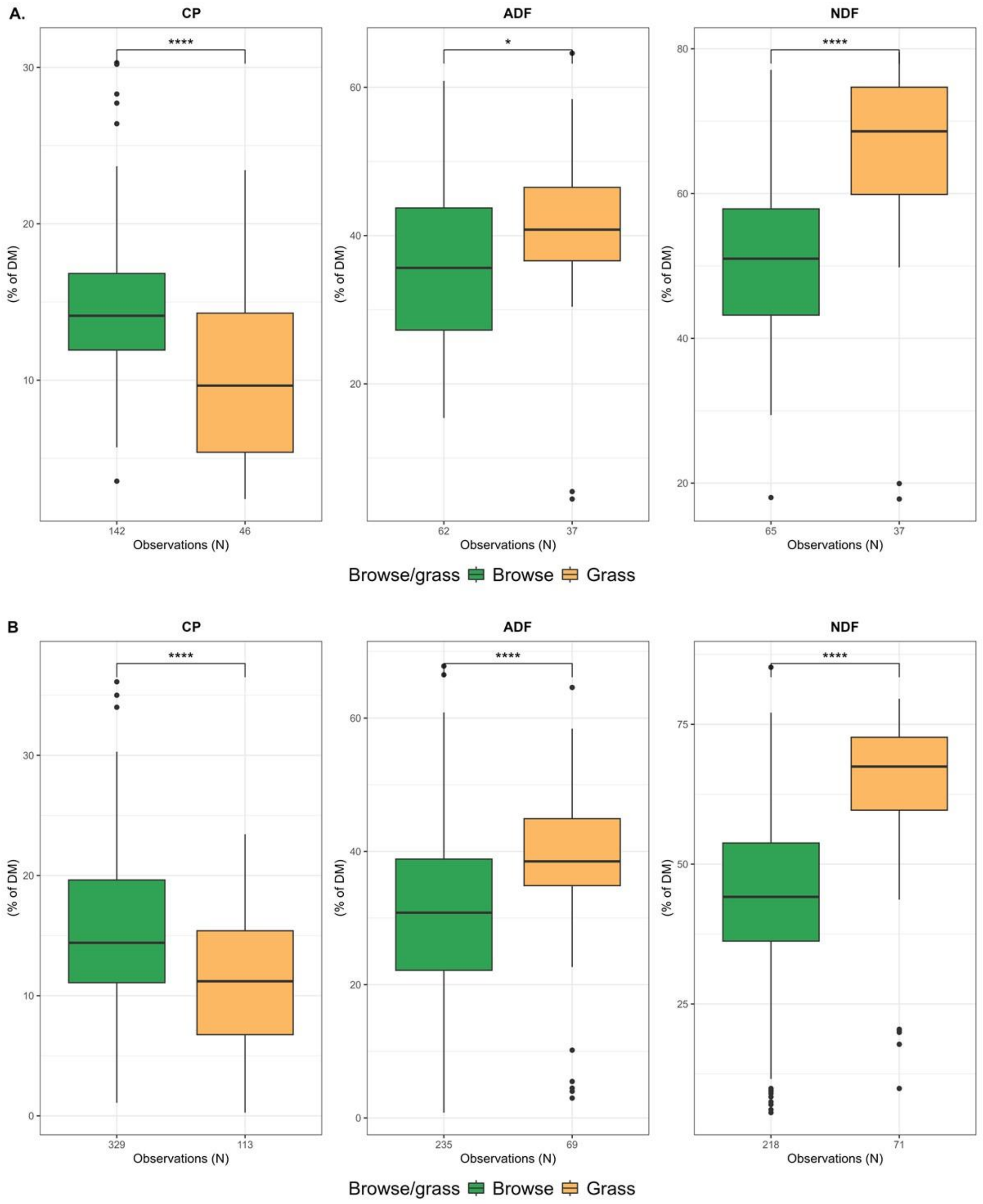


**Figure S2**: Crude protein, acid detergent fibre and neutral detergent fibre content in browse and grasses consumed by Asian elephants in dataset from **A)** Asia only and **B)** global extent. Broadly, browse food plants have higher crude protein and more digestible fibre than grasses. Note the higher CP in browse is largely because of the data being dominated by leafy browse (see Figure 2 in main text and next supplement figure).

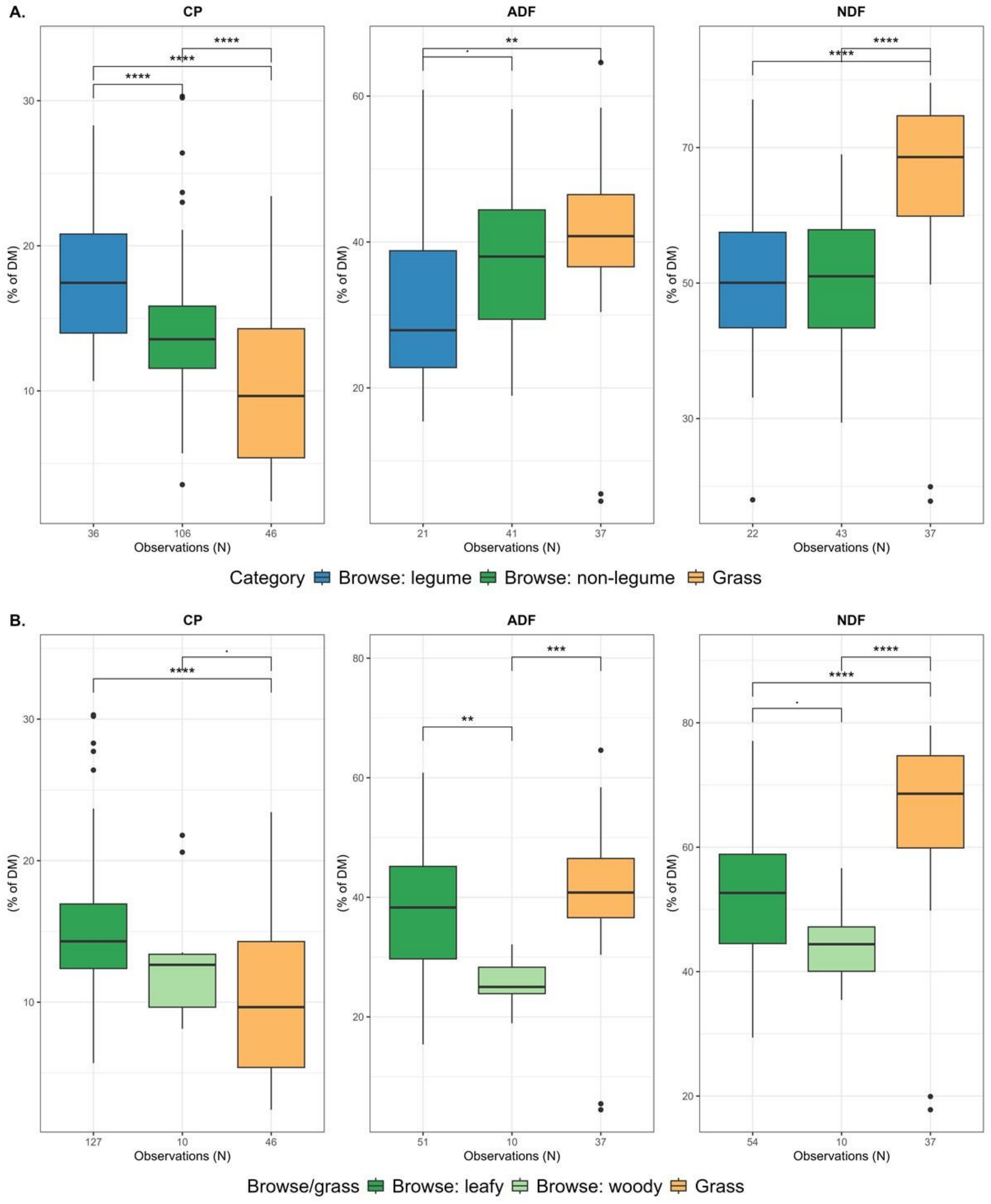


**Figure S3**. Crude protein, acid detergent fibre and neutral detergent fibre in the dataset from only Asia, for two subcategorizations **A)** legume browse, non-legume browse and grass, and **B)** leafy browse, woody browse and grass. Results from the global extent dataset are presented in the main text Figure 2.

### SUPPLEMENTARY TABLES

**Table S1**: Variance partitioning (Type III SS obtained from anova) of the fixed effects of $\delta^{13}C$ (grass-browse composition), season and their interaction in the LMMs of: **a)** N%, and **b)** C:N (log transformed) in elephant fecal samples. Estimates from the model are reported in Table S3.

| **Effect** | **SS** | **MS** | ***df*₁** | ***df*₂** | ***F*** | ***P*** |
|---|---|---|---|---|---|---|
| **(a) N%** | | | | | | |
| $\delta^{13}C$ | 0.03 | 0.03 | 1 | 83.19 | 1.471 | 0.229 |
| Season | 0.83 | 0.42 | 2 | 71.04 | 22.894 | **0.000** |
| $\delta^{13}C$ × Season | 0.10 | 0.05 | 2 | 82.95 | 2.727 | 0.071 |
| **(b) log (C:N)** | | | | | | |
| $\delta^{13}C$ | 0.12 | 0.12 | 1 | 78.99 | 4.601 | **0.035** |
| Season | 2.02 | 1.01 | 2 | 62.04 | 37.601 | **0.000** |
| $\delta^{13}C$ × Season | 0.21 | 0.10 | 2 | 77.55 | 3.841 | **0.026** |

**Table S2**: Results from the LMM showing the fixed effects of $\delta^{13}C$ (grass-browse composition, higher $\delta^{13}C$ indicates lower browsing and higher grazing), season and their interaction on **a)** N%, and **b)** C:N ratio (log transformed) in elephant fecal samples. Dry-onset is the reference season. Since $\delta^{13}C = 0$ is unrealistic for plants, the intercept values of N% and C:N were modelled for the minimum value of $\delta^{13}C$ in the dataset (-29.34).

| **Parameter** | **Estimate** | **S.E.** | **df** | **t(87)** | ***P*** |
|---|---|---|---|---|---|
| **(a) N%** | | | | | |
| Intercept | 0.90 | 0.05 | 59.67 | 17.089 | **0.000** |
| $\delta^{13}C$ | 0.03 | 0.01 | 81.45 | 3.157 | **0.002** |
| Season (dry-peak) | -0.23 | 0.09 | 78.943 | -2.713 | **0.008** |
| Season (wet) | 0.28 | 0.07 | 58.28 | 4.180 | **0.000** |
| $\delta^{13}C$ × Season (dry-peak) | -0.02 | 0.02 | 82.136 | -0.955 | 0.342 |
| $\delta^{13}C$ × Season (wet) | -0.05 | 0.02 | 83.52 | -2.313 | **0.023** |
| **(b) C:N (log transformed)** | | | | | |
| Intercept | 3.76 | 0.06 | 49.07 | 68.242 | **0.000** |
| $\delta^{13}C$ | -0.04 | 0.01 | 76.35 | -3.421 | **0.001** |
| Season (dry-peak) | 0.41 | 0.09 | 72.364 | 4.437 | **0.000** |
| Season (wet) | -0.32 | 0.07 | 47.54 | -4.556 | **0.000** |
| $\delta^{13}C$ × Season (dry-peak) | 0.00 | 0.02 | 71.543 | -0.267 | 0.791 |
| $\delta^{13}C$ × Season (wet) | 0.06 | 0.02 | 83.36 | 2.576 | **0.012** |

**Table S3**. Nutritional values of elephant food categories in the dataset from only Asia. **a)** Browse and grass, **b)** Legume browse, non-legume browse and grass, **c)** Leafy browse, woody browse and grass (values of fruit&seed browse are also reported here but were excluded from the main analyses).

**a). Browse vs. grass broad comparison**

| Category | variable | N | mean | sd |
|---|---|---|---|---|
| Browse | NDF | 65 | 50.2 | 11 |
| Grass | NDF | 37 | 65.1 | 13.7 |
| Browse | ADF | 62 | 35.9 | 11.6 |
| Grass | ADF | 37 | 41.6 | 12.3 |
| Browse | CP | 142 | 14.9 | 4.7 |
| Grass | CP | 46 | 10.1 | 5.3 |

**b). Legume browse vs. non-legume browse vs. grass**

| | variable | N | mean | sd |
|---|---|---|---|---|
| Browse: non-legume | NDF | 43 | 50.2 | 10.1 |
| Browse: legume | NDF | 22 | 50.2 | 12.8 |
| Grass | NDF | 37 | 65.1 | 13.7 |
| Browse: non-legume | ADF | 41 | 37.8 | 10.2 |
| Browse: legume | ADF | 21 | 32.2 | 13.4 |
| Grass | ADF | 37 | 41.6 | 12.3 |
| Browse: non-legume | CP | 106 | 13.9 | 4.3 |
| Browse: legume | CP | 36 | 17.7 | 4.6 |
| Grass | CP | 46 | 10.1 | 5.3 |

**c). Leafy browse vs. woody browse vs. grass**

| | variable | N | mean | sd |
|---|---|---|---|---|
| Browse: leafy | NDF | 54 | 51.9 | 10.5 |
| Grass | NDF | 37 | 65.1 | 13.7 |
| Browse: woody | NDF | 10 | 44.4 | 6.2 |
| Fruit&seed browse | NDF | 1 | 18 | NA |
| Browse: leafy | ADF | 51 | 38.3 | 11.3 |
| Grass | ADF | 37 | 41.6 | 12.3 |
| Browse: woody | ADF | 10 | 25.9 | 4 |
| Fruit&seed browse | ADF | 1 | 15.5 | NA |
| Browse: leafy | CP | 127 | 15.1 | 4.6 |
| Grass | CP | 46 | 10.1 | 5.3 |
| Fruit&seed browse | CP | 5 | 12.2 | 7.6 |
| Browse: woody | CP | 10 | 13.1 | 4.6 |

**Table S4**. Nutritional values of elephant food categories in the dataset with global extent. **a)** Browse and grass, **b)** Legume browse, non-legume browse and grass, **c)** Leafy browse, woody browse and grass (values of fruit&seed browse are also reported here but were excluded from the main analyses).

**a. Grass vs. browse broad comparison**

| gb | variable | N | mean | sd |
|---|---|---|---|---|
| Browse | NDF | 218 | 43.6 | 15.3 |
| Grass | NDF | 71 | 63.1 | 15.2 |
| Browse | ADF | 235 | 30.6 | 13.7 |
| Grass | ADF | 69 | 38.1 | 12.4 |
| Browse | CP | 329 | 15.4 | 6.5 |
| Grass | CP | 113 | 11.6 | 5.6 |

**b. Legume browse vs. non-legume browse vs. grass**

| | variable | N | mean | sd |
|---|---|---|---|---|
| Browse: non-legume | NDF | 161 | 42.8 | 15.6 |
| Browse: legume | NDF | 57 | 45.8 | 14.2 |
| Grass | NDF | 71 | 63.1 | 15.2 |
| Browse: non-legume | ADF | 176 | 30.4 | 14.3 |
| Browse: legume | ADF | 59 | 31.4 | 11.6 |
| Grass | ADF | 69 | 38.1 | 12.4 |
| Browse: non-legume | CP | 245 | 14.3 | 6.5 |
| Browse: legume | CP | 84 | 18.5 | 5.4 |
| Grass | CP | 113 | 11.6 | 5.6 |

**c. Leafy browse vs. woody browse vs. grass**

| | variable | N | mean | sd |
|---|---|---|---|---|
| Browse: leafy | NDF | 148 | 44.8 | 13.7 |
| Browse: woody | NDF | 37 | 44.7 | 13.3 |
| Fruit&seed browse | NDF | 33 | 36.7 | 21.4 |
| Grass | NDF | 71 | 63.1 | 15.2 |
| Browse: leafy | ADF | 162 | 31.2 | 12.3 |
| Browse: woody | ADF | 38 | 30.5 | 12.3 |
| Fruit&seed browse | ADF | 35 | 28.2 | 19.9 |
| Grass | ADF | 69 | 38.1 | 12.4 |
| Browse: leafy | CP | 250 | 16.7 | 5.8 |
| Browse: woody | CP | 39 | 12.9 | 6.6 |
| Fruit&seed browse | CP | 40 | 9.7 | 6.8 |
| Grass | CP | 113 | 11.6 | 5.6 |

**SI References**


1. Ahrestani, F. S., Heitkönig, I. M. A., & Prins, H. H. T. (2011). Herbaceous production in South India—Limiting factors and implications for large herbivores. Plant and Soil, 349(1–2), 319–330. https://doi.org/10.1007/s11104-011-0876-x
2. Ahrestani, F. S., Heitkönig, I. M. A., & Prins, H. H. T. (2012). Diet and habitat-niche relationships within an assemblage of large herbivores in a seasonal tropical forest. Journal of Tropical Ecology, 28(4), 385–394. https://doi.org/10.1017/S0266467412000302
3. Baskaran, N., Balasubramanian, M., Swaminathan, S., & Desai, A. A. (2010). Feeding ecology of the Asian elephant Elephas maximus Linnaeus in the Nilgiri Biosphere Reserve, southern India. Journal of the Bombay Natural History Society, 107(1), 3–13.
4. Sukumar, R. (1990). Ecology of the Asian Elephant in Southern India. II. Feeding Habits and Crop Raiding Patterns. Journal of Tropical Ecology, 6(1), 33–53.
5. Sukumar, R., & Ramesh, R. (1995). Elephant foraging: Is browse or grass more important? In Daniel, J.C. & H.S. Datye (Eds): A Week with Elephants. (pp. 368–374). Bombay Natural History Society, Oxford University Press.
6. Sukumar, R. (2003). The living elephants: Evolutionary ecology, behavior, and conservation. Oxford University Press.
